# *Candida albicans* Colonization Modulates Murine Ethanol Consumption and Behavioral Responses Through Elevation of Serum Prostaglandin E_2_ and Impact on the Striatal Dopamine System

**DOI:** 10.1101/2025.02.25.640044

**Authors:** Andrew W. Day, Jeyra Perez-Lozada, Alyssa DiLeo, Katrina Blandino, Jamie Maguire, Carol A. Kumamoto

**Author notes:** Correspondence to Carol A. Kumamoto, Department of Molecular Biology and Microbiology, Tufts University, 150 Harrison Ave., Boston, MA, 02111, USA.

## Abstract

*Candida albicans* is a commensal yeast that is a common component of the gastrointestinal (GI) microbiome of humans. *C. albicans* has been shown to bloom in the GI tract of individuals with alcohol use disorder (AUD) and can promote and increase the severity of alcoholic liver disease (ALD). However, the effects of *C. albicans* blooms on the host in the context of AUD or AUD-related phenotypes, such as ethanol preference, have been unstudied. In this work, we report a reduction in ethanol consumption and preference in mice colonized with *C. albicans*. *C. albicans-*colonized mice exhibited elevated levels of serum PGE_2_ and reduced ethanol preference was reversed by injection with antagonists of PGE_2_ receptors. Further, injection of mice with a PGE_2_ derivative decreased their ethanol preference. These results show that PGE_2_ acting on its receptors EP1 and EP2 drives reduced ethanol preference in *C. albicans-*colonized mice. We also showed altered transcription of dopamine receptors in the dorsal striatum of *C. albicans-*colonized mice and more rapid acquisition of ethanol conditioned taste aversion, suggesting alterations to reinforcement or aversion learning. Finally, *C. albicans*-colonized mice were more susceptible to ethanol-induced motor coordination impairment showing significant alterations to the behavioral effects of ethanol. This study identifies a member of the fungal microbiome that alters ethanol preference and demonstrates a role for PGE_2_ signaling in these phenotypes.

**Importance:** *Candida albicans* is a commensal yeast that is found in the gut of most individuals. *C. albicans* has been shown to contribute to alcoholic liver disease. Outside of this, the impact of intestinal fungi on alcohol-use disorder (AUD) had been unstudied. As AUD is a complex disorder characterized by high relapse rates, and there are only 3 FDA-approved therapies for the maintenance of abstinence, it is important to study novel AUD contributors to find new therapeutic targets. Here we show that an intestinal fungus, *C. albicans*, can alter mammalian ethanol consumption through an immune modulator, prostaglandin E_2_. The results highlight novel contributors to AUD-related phenotypes and further implicate the gut-brain axis in AUD. Future studies could lead to new therapeutic avenues for the treatment of AUD.

## Introduction

*Candida albicans* is a commensal fungus that is found in the gastrointestinal (GI) tract of roughly 65% of individuals (1). *C. albicans* is an opportunistic pathogen that can cause oral thrush and life-threatening bloodstream infections (2,3). Because of the severity of these infections, *C. albicans* pathogenesis has been extensively investigated. However, prior to infections, *C. albicans* lives in the gut and can breach the GI tract to infect the host (4–6).

One host setting associated with enrichment for *C. albicans* in the GI tract is alcohol use disorder (AUD), a condition in which individuals have lost the ability to control or stop their alcohol consumption (7). Estimates suggest roughly 2.3 billion people in the world consume alcohol and over 5% of the world’s adult population has an AUD (8). *C. albicans* is enriched in the fecal microbiome in individuals with AUD (9–11) and contributes to the progression and severity of ALD (9,10). Immune responses to intestinal fungi correlate with a reduced 5-year survival in those with ALD (9), showing that changes in abundance of intestinal fungi have vital consequences to the host.

The gut microbiota can also affect host behavior via effects on the gut-brain axis (GBA). The GBA is defined as the bidirectional communication between the GI tract and the brain (12,13). The GBA has been implicated in many disorders ranging from autism to addiction (14–17). Initial work characterizing the GBA in AUD found dysbiosis in the GI tract can contribute to depression and craving (18). Specific bacteria have been found that contribute to the control of factors crucial to alcohol craving (19,20). Further, fecal microbiota transplants in mice and humans can modulate ethanol preference and craving respectively (21,22). These findings support the model that changes to the microbiome in AUD can regulate ethanol consumption.

As a mechanism linking microbiome changes to the GBA in AUD, we investigated the role of prostaglandin E2 (PGE_2_), an eicosanoid compound synthesized from arachidonic acid. A previous study showed that treatment with PGE_2_ reduced ethanol consumption in rats (23). Serum PGE_2_ levels have previously been shown to be higher in antibiotic-treated mice with GI colonization by *C. albicans* (24). Further, we previously showed that *C. albicans* colonization in mice dysregulated endocannabinoids (25), another set of compounds that utilize arachidonic acid as a precursor (26). The arachidonic acid-derived host lipidome is thus altered in *C. albicans-*colonized mice. We therefore tested for a role for PGE_2_ in reducing ethanol consumption in *C. albicans-*colonized mice.

In this study, we show that *C. albicans* GI colonization in mice reduced their ethanol consumption. *C. albicans*-colonized mice had elevated serum PGE_2_ compared to mock-colonized mice, and injection of uncolonized mice with dimethyl-PGE_2_ reduced ethanol consumption. The reduction in ethanol consumption in *C. albicans-*colonized mice was reversed by antagonizing receptors for PGE_2_*. C. albicans*-colonized mice exhibited altered expression of dopamine receptors in the dorsal striatum, a more rapid acquisition of ethanol-induced conditioned taste aversion, and were more sensitive to ethanol-induced motor coordination deficits which could be reversed by PGE_2_ receptor antagonist treatment. These results show for the first time that changes to the fungal microbiome can alter ethanol preference and ethanol-induced behavior in mice.

## Results

### *C. albicans* colonization in female mice reduces ethanol consumption

Mice were orally inoculated with *C. albicans* as described (25).We previously showed that inoculation did not result in significant changes in bacterial microbiota composition and did not evoke significant inflammation (25). Colonized and mock-colonized mice were subjected to a continuous access two-bottle choice experiment where mice were given access to two sipper tubes containing water or 15% ethanol simultaneously as described in Materials and Methods. Ethanol consumption and ethanol preference showed reductions on both days of the protocol in *C. albicans*-colonized mice compared to mock-colonized mice (mock) (**Figures 1A, B**). *C. albicans*-colonized mice consumed slightly less total liquid than mock on day 1 and the same amount of liquid on day 2. Both groups drank less total liquid than mice only given access to water (H2O-only) (**Figure 1C**). Colony forming units/gram of fecal pellets (FP) or cecum contents (CC) demonstrated the presence of *C. albicans* in the GI tract of inoculated mice throughout the experiment **(Figure 1D**). From these results, we concluded that *C. albicans* colonization reduced ethanol consumption in female mice.

**Figure 1:**
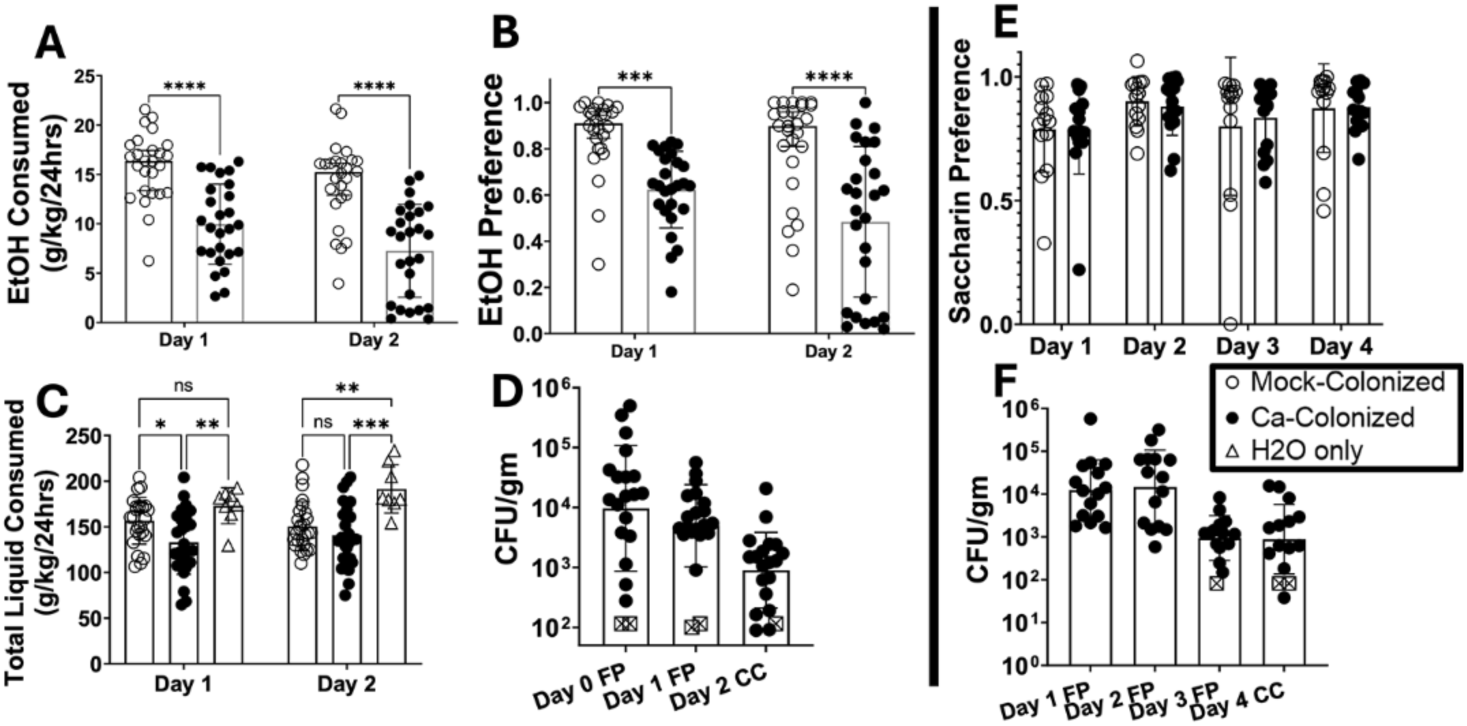
C. albicans-colonized mice consume significantly less ethanol and have lower ethanol preference than mock-colonized mice Single housed female C57BL/6 mice were orally inoculated with *C. albicans* strain CKY101 or PBS. After inoculation, mice were subjected to a 2-bottle choice experiment, with one bottle of 15% ethanol (vol/vol) and one bottle of water. Liquid consumption was measured by weighing the bottles daily. **(A)** Ethanol consumption (gm ethanol/kg body weight) on the first and second day of access to ethanol. Mock-colonized (open circles), *C. albicans-*colonized (black circles). **(B)** Ethanol preference (volume of ethanol solution consumed/total volume of liquid consumed). Mock-colonized (open circles), *C. albicans*-colonized (black circles). **(C)** Total liquid consumed per day (gm total liquid/kg body weight). Mock or *C. albicans*-colonized mice (open and black circles respectively) and mice that were only given access to water (H2O only, triangles). **(D)** CFU/gram of fecal pellet on each day of collection. FP, fecal pellets; CC, cecum contents. Black circles show individual mice and open squares show values below the limit of detection. Geometric means are shown by bars and geometric standard deviation is shown by error bars. **(E)** Saccharin preference (volume of saccharin solution/total volume of liquid consumed). **(F)** CFU/gm fecal pellets from the saccharin preference experiment. Symbols as in panel D. **(A-C,E)** Boxes represent mean values, error bars show standard deviation, and each symbol represents an individual mouse. A two-way ANOVA corrected for repeated measures was performed for statistics (* p=0.0144; ** p<0.0035; *** p<0.0003; **** p<0.0001).

Saccharin preference was tested to determine whether *C. albicans* colonized mice showed alterations in preference to other rewarding substances (27). Saccharin preference was measured in a similar continuous access two-bottle choice experiment as described in Materials and Methods. Saccharin preference per day showed no effect of *C. albicans* colonization (**Figure 1E)**. CFU/gram of FP or CC showed colonization throughout the experiment (**Figure 1F)**. These results showed that *C. albicans* colonization did not exert broad effects on consumption of all rewarding substances. Rather, the effects were more selective.

### *C. albicans*-colonized mice have elevated serum PGE_2_ and PGE_2_ injection reduces murine ethanol preference

Levels of PGE_2_ were measured in serum obtained from *C. albicans*-colonized or mock-colonized mice following two days of ethanol consumption. Since PGE_2_ is not a stable molecule, concentrations of a metabolite of PGE_2_, 13,14-dihydro-15-keto PGE_2_ (PGE-m), commonly utilized as a marker for PGE_2_ (28), were assayed by ELISA. The assay showed that *C. albicans*-colonized mice had elevated serum PGE-m levels compared to H_2_O-only or mock groups (**Figure 2A**).

**Figure 2.**
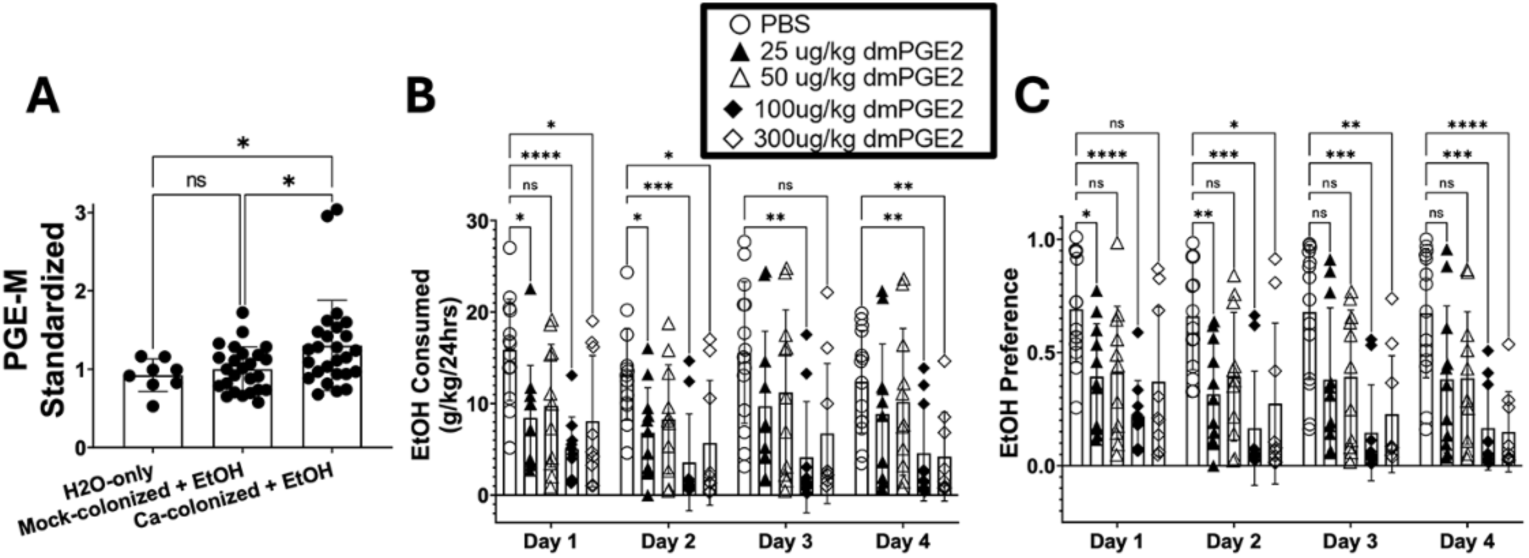
C. albicans-colonized mice have elevated serum PGE_2_ and injection of dmPGE_2_ reduced ethanol consumption. Single housed female C57BL/6 mice were orally inoculated with *C. albicans* strain CKY101 or PBS and subjected to the 2-bottle choice experiment as in Figure 1. On day 2, mice were euthanized, and serum was collected. **(A)** PGE-metabolite concentration, standardized to the average in mock-colonized mice within each experiment. Mice only given access to H_2_O (left bar), mock-colonized mice that consumed ethanol (middle bar), *C. albicans*-colonized mice that consumed ethanol (right bar). Bars show mean, error bars show standard deviation, each symbol indicates an individual mouse. One-way ANOVA was performed for statistical analysis (*, p<0.0354) **(B)** Single housed female C57BL/6 mice were injected subcutaneously once per day with dmPGE_2_ (concentrations as shown) or vehicle and subjected to the 2-bottle choice experiment as in Figure 1. On day 4, mice were euthanized, and brains were collected. Ethanol consumed per day (gm ethanol/kg body weight) is shown. **(C)** Daily ethanol preference (volume ethanol solution consumed/total volume liquid consumed) is shown. **(C and D)** Mean and standard deviation are shown. Each symbol represents one mouse. Two-way ANOVA test corrected for repeated measures was used for statistical significance. ns p<0.0939; * p<0.0499; ** p<0.0082; *** p<0.0011; ****, p<0.0001.

To test the effect of elevated PGE_2_ on ethanol consumption, a 4 day, two-bottle choice experiment was performed in which mice were injected with 16,16-dimethyl PGE_2_ (dm-PGE_2_), a commonly-used, stable derivative of PGE_2_ (29). Mice received either vehicle or one of four different concentrations of dm-PGE_2_ (25, 50, 100, 300 ug/kg) by subcutaneous injection each day. Lower concentrations (25 or 50 ug/kg) of dm-PGE_2_ led to low-level decreases in ethanol preference and consumption, and higher concentrations led to larger decreases compared to PBS-injected mice (**Figures 2B, C**). Thus, dm-PGE_2_ injection in mice decreased ethanol consumption in a dose-dependent manner, supporting the model that elevation of endogenous PGE_2_ results in decreased ethanol consumption.

### EP Receptor Antagonism Increases Ethanol Consumption and Preference to Mock-Colonized Levels

Based on these results, we hypothesized that PGE_2_ is a key driver of reduced ethanol preference in *C. albicans*-colonized mice. To test this hypothesis, female mice were orally inoculated with *C. albicans* or mock-colonized with PBS and injected with 2 g/kg each of an EP1 antagonist (SC-51089, Cayman Chemical) and an EP2 antagonist (TG6-10-1, Cayman Chemical) subcutaneously each day of the 2 day two-bottle choice experiment. Mock-colonized mice injected with vehicle or antagonist showed no difference in ethanol consumption or preference (**Figure 3A,B**). Mice that were colonized with *C. albicans* and injected with vehicle showed significantly lower ethanol consumption and preference compared to other groups. Those injected with antagonists showed significantly higher ethanol consumption and preference on day 1 and a trend towards an increase in ethanol consumption on day 2. Some reductions in total liquid consumed in *C. albicans*-colonized, vehicle-injected mice were observed (**Figure 3C**). Mice had stable colonization throughout the experiment (**Figure 3D**).

**Figure 3.**
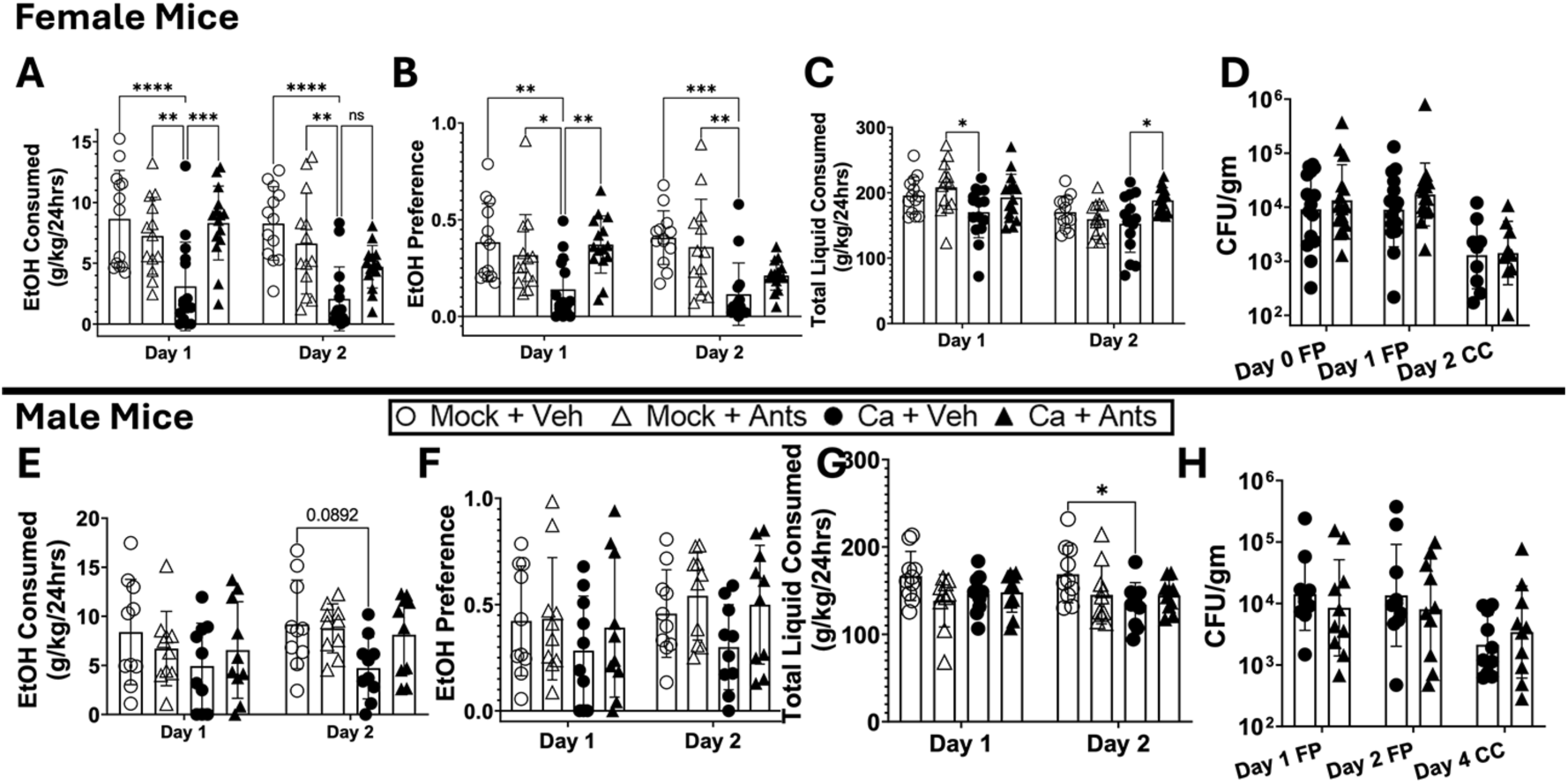
EP receptor antagonism elevates C. albicans-colonized mouse ethanol consumption to mock-colonized levels in female and male mice. Single housed female C57BL/6 mice **(A-D)** or male C57BL/6 mice **(E-H)** were orally inoculated with *C. albicans* strain CKY101 or PBS, injected subcutaneously with EP1 antagonist (SC-51089, 2 g/kg) and EP2 antagonist (TG6-10-1, 2 g/kg) or vehicle and subjected to the 2-bottle choice experiment as in Figure 1. Females were euthanized on day 2, males on day 4. **(A, E)** ethanol consumption **(B, F)** ethanol preference **(C, G)** total liquid consumed per day **(A-C, E-G)** mock-colonized mice with vehicle (open circles) or antagonists (open triangles), *C. albicans*-colonized mice with vehicle (black circles) or antagonists (black triangles). Boxes represent mean values, error bars show standard deviation, and each symbol represents an individual mouse. Two-way ANOVA corrected for repeated measures was performed for statistics. ns p=0.1182; * p<0.0368; ** p<0.0054; *** p=0.0001; **** p<0.0001. **(D, H)** shows CFU/gram of fecal pellet on each day of collection. FP, fecal pellets; CC, cecum contents. Black circles (*C. albicans* with vehicle) or black triangles (*C. albicans* with antagonists) show individual mice. Geometric means are shown by bars, and geometric standard deviation is shown by error bars. Two-way ANOVA performed for statistics.

We also tested whether male mice had a reduction in ethanol consumption following *C. albicans* colonization and whether this phenotype could be reversed by EP antagonists. A trend towards reduction in ethanol consumption by the vehicle injected, *C. albicans*-colonized mice was observed on day 2 of ethanol consumption, and total liquid-consumed was also reduced (**Figures 3E,F,G**). These mice had stable colonization throughout the experiment (**Figure 3H)**. These results support the model that activation of EP1/EP2 receptors in *C. albicans*-colonized mice decreases their ethanol consumption and preference. Further experiments were conducted with females only since females showed strong effects of colonization.

### Increased Serum PGE_2_ in *C. albicans*-colonized Mice Correlates with Gene Expression in the Brain

We next tested the hypothesis that *C. albicans* gut colonization and elevation of PGE_2_ concentration would result in transcriptomic changes in the brain. PGE_2_ binds to its cognate receptors encoded by four different genes, *Ep1*, *Ep2*, *Ep3*, and *Ep4*(30). PGE_2_ has been shown to upregulate expression of *Ep* receptor genes in a dose dependent manner in neuronal cell lines and various tissues in the body(31,32). We decided to focus on a region of the brain involved in habit formation, the dorsal striatum (DS) (33), as well as other addiction-implicated regions such as the prefrontal cortex (PFC) and nucleus accumbens (NAc)(34).

To test the hypothesis, RT-qPCR from DS-cDNA was performed and expression of *Ep1* and *Ep2* relative mock-colonized from the same experimental trial was measured. There were no significant differences in *Ep* receptor expression between groups in the DS, PFC, or NAc (**Figure S1**). However, correlations between PGE-m concentration in the serum and relative expression of *Ep1* or *Ep2* in the same *C. albicans*-colonized mouse were observed. Gene expression relative to the average for mock-colonized was plotted against the concentration of PGE-m (also standardized to mock-colonized). **Figure 4A,B** demonstrates that mice with higher concentrations of serum PGE_2_ exhibited higher expression of *Ep1* and *Ep2.* In mock-colonized mice, there were no significant correlations (**Figure 4C,D).** We observed similar significant correlations of *Ep1* expression and PGE-m concentration in the PFC and NAc (**Figure S2**). Elevated PGE_2_ levels were thus associated with higher expression of *Ep* receptors in multiple regions of the brain.

**Figure 4.**
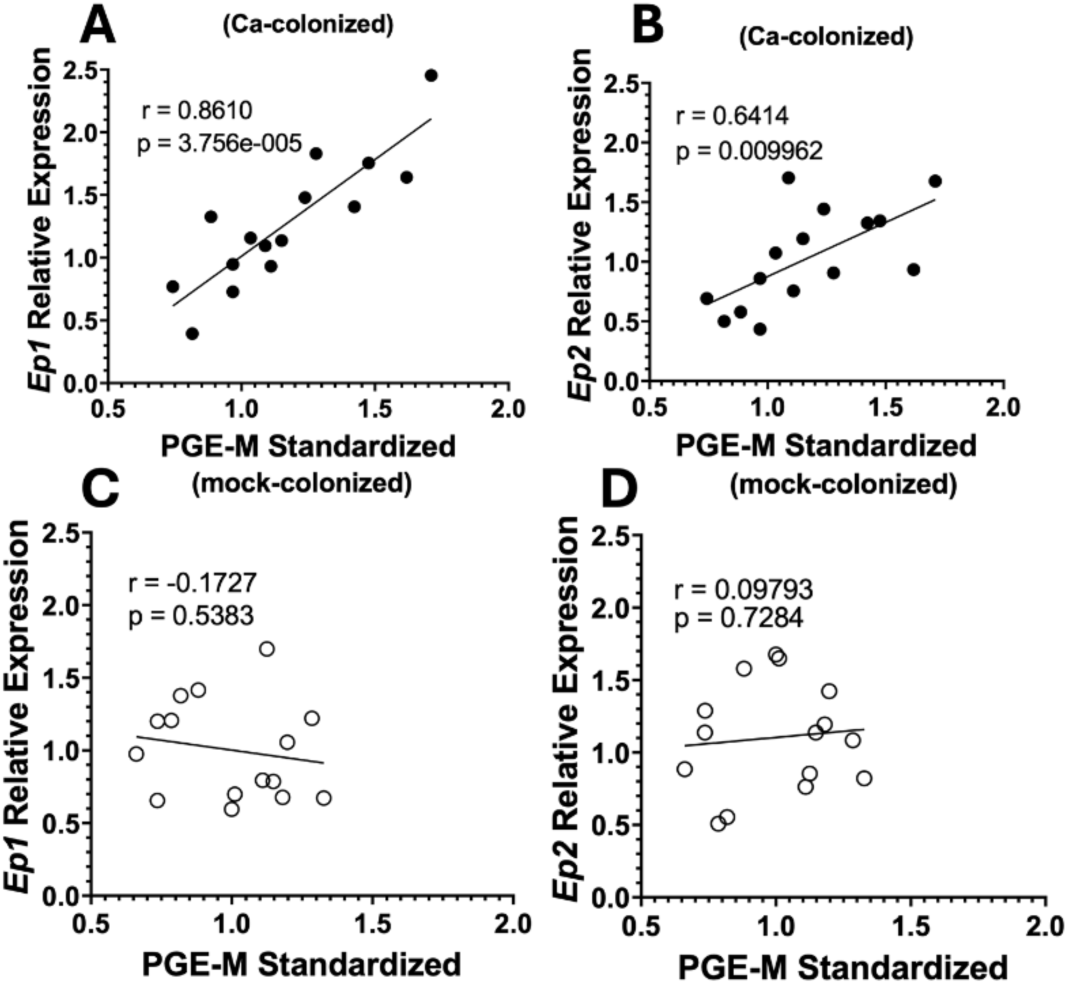
Serum PGE-metabolite correlates with EP Receptor expression in the dorsal striatum in C. albicans-colonized mice. Single housed female C57BL/6 mice were orally inoculated with *C. albicans* strain CKY101 or PBS and subjected to the 2-bottle choice experiment as in Figure 1. On day 2, mice were euthanized, and brain and serum were collected. Expression of *Ep* genes relative to average expression in mock-colonized mice from the same experimental trial, plotted as a function of concentration of PGE-metabolite standardized to the average in mock-colonized mice in the same experimental trial. Each symbol represents an individual mouse. **(A,B)** C. albicans-colonized mice (black circles) **(C,D)** mock-colonized mice (open circles). **(A,C)** Ep1 **(B,D)** Ep2. Pearson correlations were used to test for significant correlations, r=correlation strength and p=statistical significance. Significant correlations were observed between *Ep1* and *Ep2* expression and concentration of PGE-M in *C. albicans-*colonized mice.

We also analyzed the expression of dopamine receptors in the DS of these mice. The dopamine system in the DS is commonly involved with learning and habit formation (35). Expression of two main classes of dopamine receptors, *Drd1* and *Drd2*, was measured (36). A trend towards a reduction in *Drd1* expression and a statistically significant reduction in *Drd2* expression in *C. albicans*-colonized mice compared to mock-colonized were observed (**Figure 5A**). Additionally, mice injected with low-dose dm-PGE_2_ (25 or 50ug/kg injections) showed reduced expression of *Drd1* and *Drd2* (**Figure 5B**). Interestingly, *Drds* showed no differences in expression in the PFC or NAc (**Figure S3**). Furthermore, other addiction-related genes showed no significant differences in expression in the DS (**Figure S4**). These results show that both *C. albicans* colonization and low-dose dm-PGE_2_ injections reduce dopamine receptor expression in the DS.

**Figure 5.**
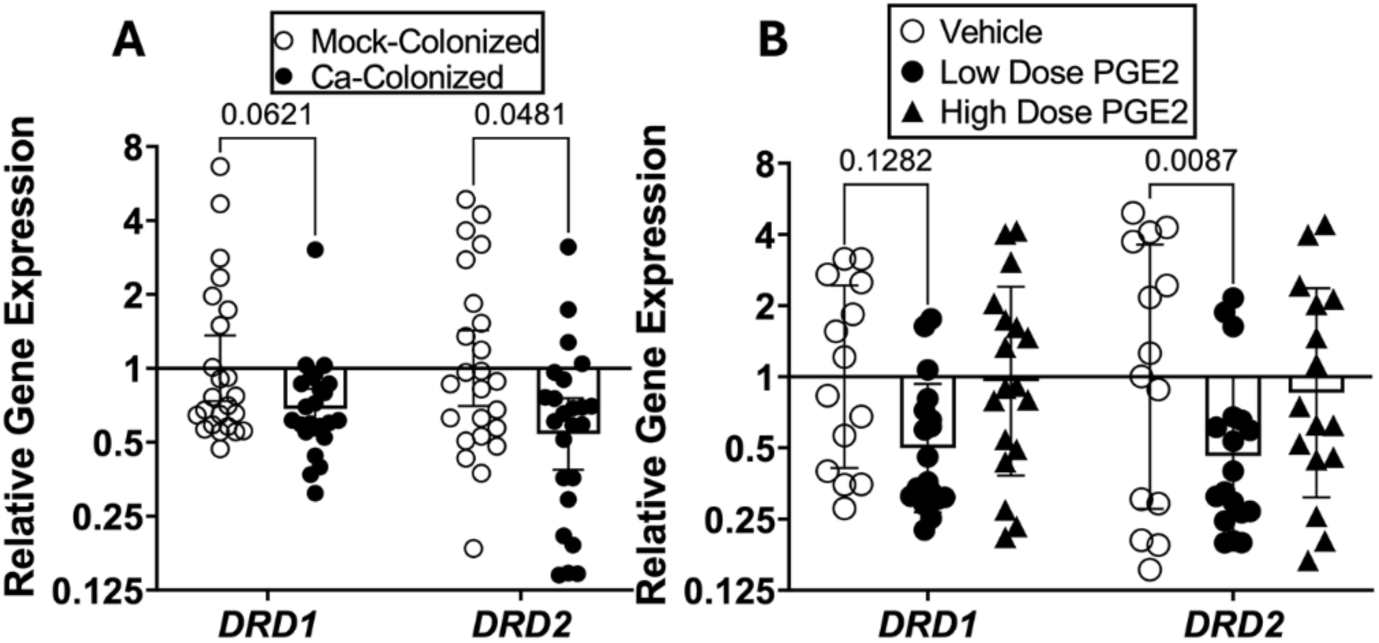
Both C. albicans colonization and low dose dmPGE2 injection result in lower Drd2 receptor expression in the Dorsal Striatum. **(A)** Single housed female C57BL/6 mice were orally inoculated with *C. albicans* strain CKY101 or PBS and subjected to the 2-bottle choice experiment as in Figure 1. On day 2, mice were euthanized, and brains were collected. *Drd* receptor expression was measured in the dorsal striatum of mock-colonized or *C. albicans*-colonized mice by real time RT qPCR using the ddCT method. **(B)** Single housed female C57BL/6 mice were injected subcutaneously each day with dmPGE_2_ (low dose = 25 or 50 ug/kg; high dose = 100 or 300 ug/kg) or vehicle and subjected to the 2 bottle choice experiment as in Figure 1. On day 4, mice were euthanized, and brains were collected. *Drd* receptor expression was measured in the DS by RT-qPCR using the ddCT method. Outliers were removed with ROUT to determine mathematical outliers (Q=0.1% to remove definitive outliers). **(A,B)** Geometric means and geometric SDs are shown and Two-way ANOVA was performed for statistical significance.

Expression of *Drd1* or *Drd2* in mock or *C. albicans*-colonized mice correlated with total ethanol consumed. In the dorsal striatum, DRD1 is involved with reward formation and DRD2 is involved with aversion formation (37). Plots of *Drd* expression and total ethanol consumed showed a significant positive correlation between *Drd1* expression and total ethanol consumption in mock-colonized mice (**Figure 6A**) and a trend towards a positive correlation between *Drd2* and total ethanol consumed (**Figure 6C**). In *C. albicans*-colonized mice, by contrast, a significant negative correlation between *Drd2* expression and total ethanol consumption was observed (**figure 6D**). No correlation between *Drd1* expression and total ethanol consumed was observed in these mice (**Figure 6B**). Thus, *C. albicans-*colonized mice that consumed lower amounts of ethanol exhibited higher expression of the aversion-associated DRD2 receptor.

**Figure 6:**
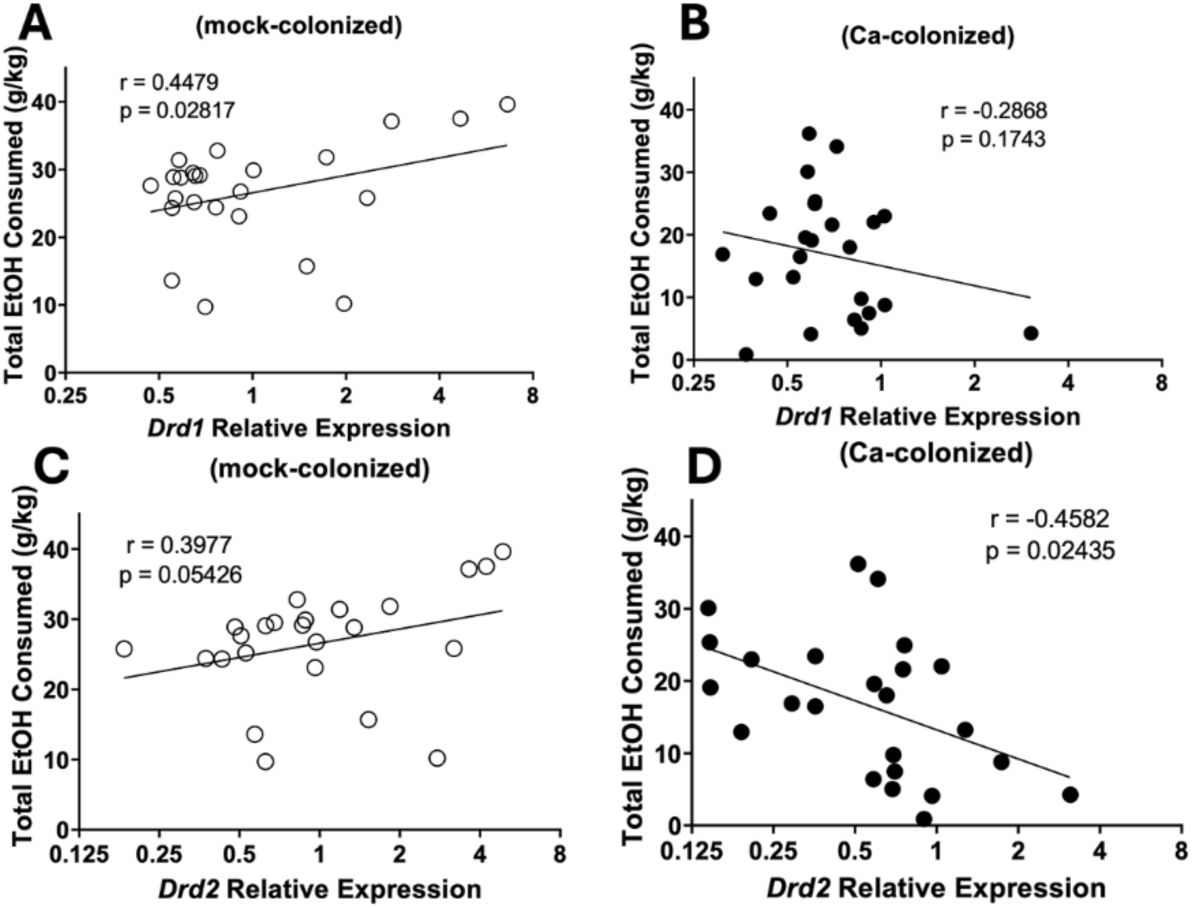
Drd Expression correlates with ethanol consumption in mock and C. albicans-colonized mice. Single housed female C57BL/6 mice were orally inoculated with *C. albicans* strain CKY101 or PBS and subjected to the 2 bottle choice experiment as in Figure 1. On day 2, mice were euthanized and brains were collected. *Drd* receptor expression was measured in the dorsal striatum of mock-colonized or *C. albicans*-colonized mice. Total ethanol consumption (gm/kg) is plotted as a function of relative *Drd* expression. Pearson correlation used to test for significant correlations, r=correlation strength and p=statistical significance. **(A, C)** mock-colonized mice. **(B, D)** *C. albicans-*colonized mice. **(A, B)** *Drd1* fold change of relative expression. **(C, D)** *Drd2* fold change of relative expression. Significant correlations between *Drd1* expression and ethanol consumption by mock-colonized mice and *Drd2* expression and ethanol consumption by *C. albicans* colonized mice were observed.

### *Candida albicans* colonization accelerates the onset of conditioned taste aversion in mice

Based on the low ethanol consumption and negative correlation between *Drd2* expression and ethanol consumption in *C. albicans*-colonized mice, we hypothesized that these mice would show enhanced ethanol-induced conditioned taste aversion (CTA). Ethanol-induced CTA is based on the principle that mice will develop an aversion to a novel tasting solution when it is paired with an ethanol injection (38,39). Mice were tested for ethanol-induced CTA as described in Materials and Methods.

The first exposure to the tastant showed the baseline consumption level, prior to injection (conditioning) with ethanol or control (sterile saline). Trials 2-4 showed consumption after conditioning, expressed as the change from initial consumption (**Figure 7A**). In trial 2, control mice showed similar or higher than baseline consumption of the tastant. Both mock-colonized and *C. albicans*-colonized mice injected with 3g/kg ethanol developed a near complete aversion to the tastant (100% reduction), which continued through trials 3 and 4. In contrast, mice given a lower concentration of ethanol (2g/kg) showed differences between groups. There was no significant aversion in the mock-colonized mice in trial 2 of this injection group and *C. albicans*-colonized mice showed a significant reduction in saline consumption compared to control mice in trial 2. On trial 3, both mock-colonized and *C. albicans*-colonized mice injected with 2g/kg ethanol showed significant aversion to the tastant. There were no differences in colonization or mouse weight throughout the experiment (**Figure 7B, S5**). *C. albicans-*colonized mice thus developed ethanol-induced CTA more rapidly.

**Figure 7:**
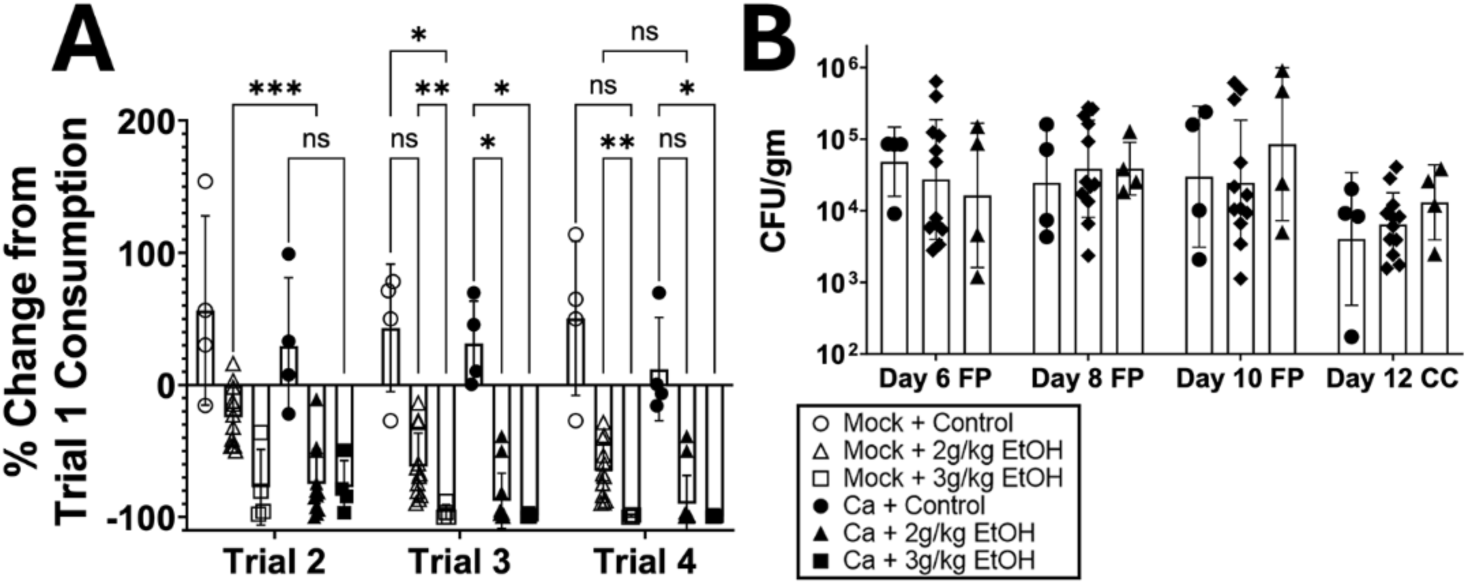
C. albicans-colonized mice develop conditioned taste aversion more rapidly than mock-colonized mice. Single housed female C57BL/6 mice were trained to drink their daily liquid in the late morning. Mice orally inoculated with *C. albicans* strain CKY101 or PBS were given one hour of access to a novel tastant, 1.2% saline solution, and then injected intraperitoneally with ethanol (2 g/kg or 3 g/kg) or control (sterile saline). Tastant consumption on the first trial (before injection with ethanol) was used as the baseline measurement and consumption on later trials was compared to trial 1. **(A)** percent change of tastant consumption relative to each individual mouse’s trial one consumption (pre-conditioned stimulus). Trial 1 corresponds to Day 6 of the experiment, Trial 2 corresponds to Day 8 of the experiment, Trial 3 corresponds to Day 10 of the experiment, and Trial 4 corresponds to Day 12. **(B)** CFU/gm of *C. albicans* from fecal pellets collected on Day 6, Day 8, and Day 10; cecum contents from Day 12. **(A,B)** Mock-colonized, control (open circles); mock-colonized + 2g/kg EtOH (open triangles); mock-colonized + 3 g/kg EtOH (open squares); *C. albicans*-colonized, control (closed circles); *C. albicans-*colonized +2 g/kg EtOH (black triangles); *C. albicans-*colonized + 3 g/kg EtOH (closed squares). Two-way ANOVA corrected for repeated measures was completed for statistical significance. ns p<0.0995; * p<0.0499; ** p<0.0076; *** p=0.0006.

### *C. albicans*-colonized mice are more susceptible to the behavioral effects of ethanol and motor coordination deficits can be restored by EP receptor antagonism

The more rapid development of ethanol-induced CTA in *C. albicans*-colonized mice suggests these mice could be more susceptible to the behavioral effects of ethanol as correlations between these phenotypes have been observed (40). Therefore, we tested whether *C. albicans*-colonized mice were more susceptible to ethanol sedation, ethanol-induced motor coordination defects, and depressant effects of ethanol (**Figure 8A**).

**Figure 8:**
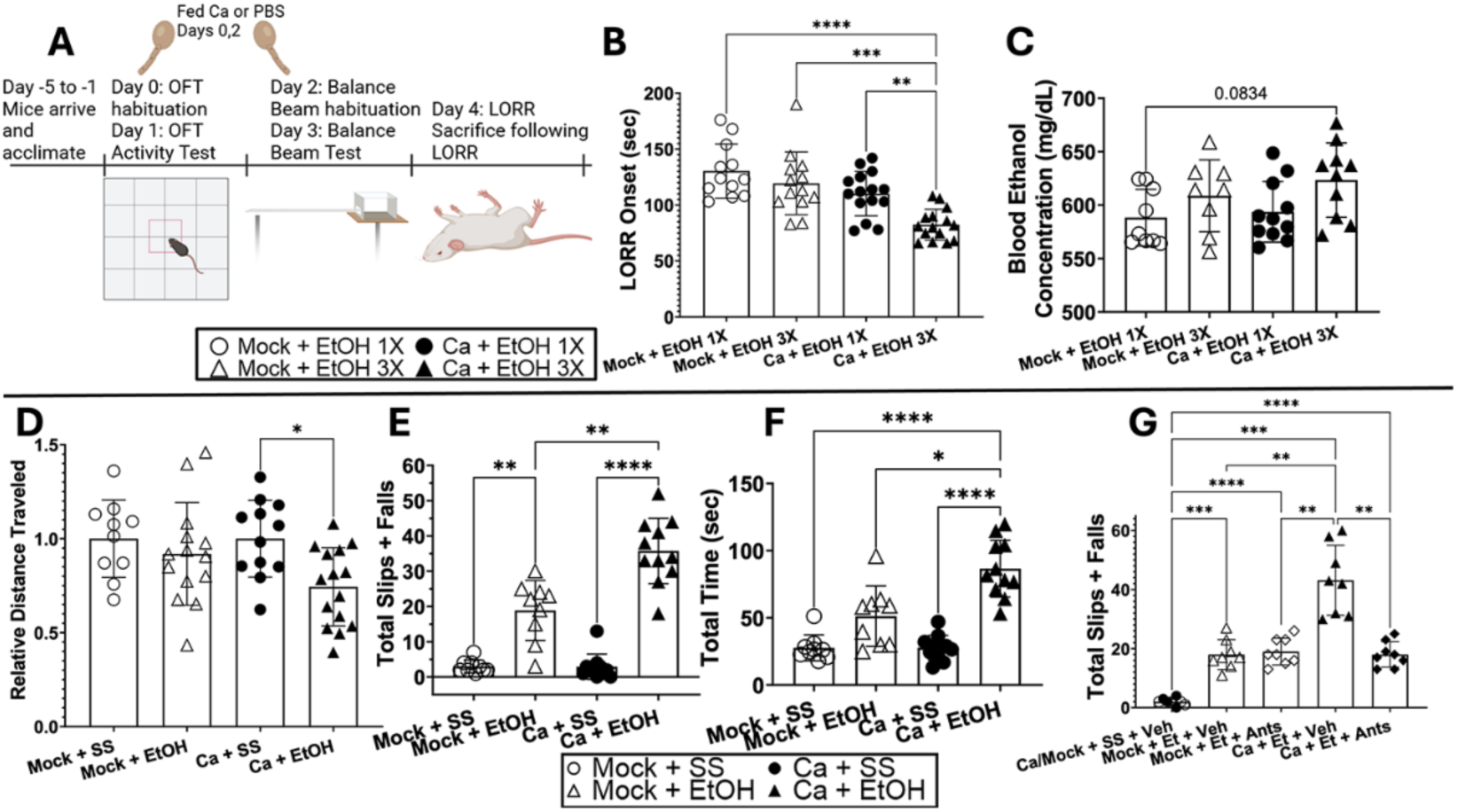
C. albicans-colonized mice are more sensitive to effects of ethanol. Single housed female C57BL/6 mice were orally inoculated with *C. albicans* strain CKY101 or PBS on days 0 and 2, and subjected to the behavioral tests illustrated in panel **(A). (B,C)** Groups are defined as Mock +1x EtOH: mock-colonized, received sterile saline ip injections on days 1 and 3 and 3.5g/kg EtOH on day 4 (open circles); Mock + 3x EtOH: mock-colonized, received 1.5g/kg ethanol ip injected on days 1 and 3 and 3.5g/kg EtOH on day 4 (open triangles); Ca + 1x EtOH: *C. albicans*-colonized, received sterile saline ip injections on day 1 and 3 and 3.5g/kg EtOH on day 4 (black circles); Ca + 3x EtOH: *C. albicans*-colonized, received 1.5g/kg ethanol ip injected on days 1 and 3 and 3.5g/kg EtOH on day 4 (black triangles). (**D-F)** Groups are defined as Mock + SS: mock-colonized, received sterile saline ip injections on days 1 and 3 (open circles); Mock + EtOH: mock-colonized, received 1.5g/kg ethanol ip injected on days 1 and 3 (open triangles); Ca + SS: *C. albicans*-colonized, received sterile saline ip injections on day 1 and 3 (black circles); Ca + EtOH: *C. albicans*-colonized, received 1.5g/kg ethanol ip injected on days 1 and 3 (black triangles). **(G)** Groups are defined as Ca/Mock + SS + Veh, fed *C. albicans* (black circles) or sterile PBS (open circles), received sterile saline and vehicle injections; Mock + Et + Veh, fed sterile PBS, received ethanol and vehicle injections; Mock + Et + Ants, fed sterile PBS, received ethanol and antagonist injections; Ca + Et + Veh, fed *C. albicans*, received ethanol and vehicle injections; Ca + Et + Ants, fed *C. albicans*, received ethanol and antagonist injections. All injections were IP, given before habituation or testing. Ethanol injections were 1.5g/kg of mouse weight and antagonists were 2g/kg of both EP1 and EP2 antagonists. **(B)** time after 3.5 g/kg ethanol injection (seconds) for the mouse to lose its righting reflex (LORR onset). **(C)** blood ethanol concentration (mg/dL) at the time of euthanization following LORR onset. **(D)** distance traveled in the open field test in a ten minute trial, shown relative to the average of the sterile saline-injected (SS) mice within the same group (mock-colonized or Ca-colonized). **(E)** total number of foot slips and “falls” during three trials of the balance beam test. A foot slip was defined as the foot slipping beneath the balance beam and a fall was defined as falling to the underside of the beam, or, on rare occasions, a fall off the beam. **(F)** cumulative time required to cross the balance beam in three trials. **(G)** total number of slips or “falls” from the EP-antagonist experiments. Mice were injected intraperitoneally with EP1 and EP2 antagonists or vehicle daily on days 0-4 approximately one hour before each behavioral test. **(B,C)** Ordinary One-Way ANOVAs were completed for statistical significance. **(D-G)** Brown-Forsythe and Welch ANOVAs were completed for statistical significance. **(B-G)** Bars show means with the standard deviation or geometric mean with geometric standard deviation. Symbols represent individual mice. * p<0.0220; ** p<0.0055; *** p<0.0003; **** p<0.0001.

The first experiment was the open field activity test (OFT), which measures how susceptible C57BL/6 mice are to the depressant effects of alcohol based on the decrease in distance traveled in the OFT following ethanol injection (1.5 g/kg) (41). Mice were injected with ethanol (dose) or control (sterile saline [SS]) and distance traveled in 10 minutes in the OFT was measured. Results are expressed relative to the SS-injected mice within each experimental trial (mock or *C. albicans*-colonized) (**Figure 8B**). Results showed a significant reduction in distance traveled by *C. albicans*-colonized, ethanol-injected mice compared to *C. albicans*-colonized, SS-injected mice. There was not a significant reduction in distance traveled by the mock-colonized, ethanol-injected mice compared to SS-injected mice. These results show that the *C. albicans*-colonized mice were more strongly affected by ethanol.

The loss of righting reflex (LORR) experiment tested the ability of mice to right themselves after injection with a sedative amount of ethanol (3.5 g/kg) (42). The time for the mice to lose their righting reflex after the injection is termed LORR onset. Mock or *C. albicans*-colonized mice that had received SS injections the previous three days, or mock mice that received ethanol the previous 3 days had no differences in LORR onset (**Figure 8C**). *C. albicans*-colonized mice that received ethanol injections on the previous 3 days showed a significantly shorter LORR onset compared to all groups. Blood ethanol concentration from serum collected at the time of loss of righting reflex showed a trend towards higher levels in these mice (**Figure 8D**). These results indicate a synergistic effect of ethanol treatment (including prior treatment) and *C. albicans*-colonization that increased the sedative effects of ethanol.

Motor coordination defects were tested using performance on the balance beam, which can measure coordination based on the number of slips and falls on the beam (43). Mice were injected with SS or ethanol and their performance was assessed. The total number of slips or falls throughout three trials (**Figure 8E**) and the total amount of time required to complete three trials were measured (**figure 8F**). Mock-colonized or *C. albicans*-colonized mice injected with SS did not slip/fall often and completed the trials quickly. Mock-colonized mice injected with ethanol showed a significant increase in slips and falls, consistent with ethanol-induced motor coordination deficits. *C. albicans*-colonized mice injected with ethanol showed significantly more slips and falls than any of the other groups and were significantly slower. Furthermore, there were no differences in colonization or mouse weight in these experiments (**Figure S6**). Thus, *C. albicans*-colonized mice showed more ethanol-induced impairment in motor coordination.

To test the hypothesis that PGE_2_ signaling contributed to the differences in mouse susceptibility to the effects of ethanol, mice were injected with the combination of EP1 and EP2 antagonists (2mg/kg each) before each behavioral test. Total slips and falls on the balance beam test following these injections was measured. *C. albicans*-colonized mice injected with ethanol and antagonists showed a significant decrease in slips and falls compared to the ethanol and vehicle group (**Figure 8G**). There was no difference between the *C. albicans*-colonized group with ethanol and antagonists compared to either of the mock + ethanol injected groups. We did not see a similar reversal of phenotypes by antagonists in either the OFT or LORR (**Figure S7**). These results showed that blocking PGE_2_ signaling in *C. albicans*-colonized mice reduced their susceptibility to ethanol-induced impairment of motor control.

Taken together, the results showed that *C. albicans*-colonized mice were more susceptible to the depressant and sedative effects of ethanol, and to ethanol-induced impairment in motor control. PGE_2_ effects on EP1 and EP2 receptors make a major contribution to the mechanism of ethanol-induced motor impairment.

## Discussion

We demonstrate that *C. albicans*-colonization in mice reduces ethanol preference and consumption. Our results with EP1/EP2 antagonists show that PGE_2_ signaling plays a significant role in modifying mouse ethanol preference. Further, injection of uncolonized mice with dm-PGE_2_ reduced ethanol consumption. There was a somewhat dose-dependent response as low concentrations of dm-PGE_2_ (25-50 ug/kg) reduced consumption moderately and higher doses of dm-PGE_2_ (100-300ug/kg) reduced ethanol preference and consumption more strongly.

Antibiotic treated mice with *C. albicans* gut colonization have been previously shown to exhibit elevated serum PGE-m levels (24). Mice treated with antibiotics but not inoculated with fungi showed a modest increase in serum PGE-m levels and this level was sufficient to promote inflammatory cell responses. *C. albicans*-colonized mice in our study, which did not use antibiotics, also showed modestly increased serum concentrations of PGE-m and PGE-m concentration correlated with ethanol preference. These results indicate that modest increases in serum PGE_2_ are associated with changes in host immune responses and behavior. PGE_2_ in circulation may be directly involved in these changes or may serve as a marker indicating higher levels of PGE_2_ in multiple organs. In fact, we saw evidence of increased PGE_2_ by correlations with *Ep* expression in all regions of the brain examined.

PGE_2_ has previously been shown to alter the permeability of the blood brain barrier (BBB) by inducing alterations in endothelial permeability or by stimulating the retraction of pericytes from the surface of the BBB (44–46). PGE_2_ has also been shown to induce vasodilation (44). Vasodilatory effects and BBB alterations could lead to higher amounts of PGE_2_ in the brain and reduce ethanol consumption. These results support the model that, as PGE_2_ in the circulation elevates above normal fluctuating ranges, BBB permeability increases, and more PGE_2_ can enter the brain, resulting in correlations between its concentration in serum and *Ep1/Ep2* receptor expression in *C. albicans*-colonized mice.

Synergy between the effects of *C. albicans* colonization and the effects of ethanol were also observed. Mice that received both ethanol and *C. albicans* were the only group with significantly reduced locomotion in the OFT. *C. albicans*-colonized mice that received multiple injections with ethanol showed significantly shorter time to LORR onset than *C. albicans*-colonized mice that received one injection. Ethanol also affects the BBB through changes in integrity and permeability (47,48). Inflammatory molecules such as LPS can also synergistically increase BBB permeability (49). The effects of ethanol on the BBB may thus synergize with the effects of PGE_2_, resulting in significant behavioral effects in colonized mice. Saccharin preference may not be reduced by *C. albicans* colonization because of the lack of an additional insult to the BBB.

PGE_2_ has previously been shown to affect the dopamine system. Dopamine is critical for habit, reward, and aversion formation (50). PGE_2_ treatment altered reward learning to morphine in mice (51,52), and has been shown to alter DRD signaling and behavior following cocaine ingestion (53). We observed that *C. albicans*-colonized mice showed altered expression of dopamine receptors in the dorsal striatum. Expression of one of the receptors, *Drd2*, was significantly decreased in *C. albicans* colonized mice and low-dose dm-PGE_2_-injected mice. Interestingly, expression in the PFC and NAc did not indicate similar changes to the dopamine system, and other genes implicated in addiction such as neuropeptide Y receptors, opioid receptors (Mu and Delta), proopiomelanocortin (*Pomc*), and *Fos* were unchanged. Significant expression differences of the Mu opioid receptor between mock and *C. albicans*-colonized mice appeared to be due to differences in ethanol consumption. These results indicate an altered regulation of the dopamine system in the DS observed in mice with low level increases in systemic PGE_2_ concentrations.

Furthermore, when *Drd* expression in the DS was plotted against total ethanol consumption, significant positive or negative correlations were observed. DRD1 and DRD2 receptors in the dorsal striatum have well defined, consistent roles. DRD1 signaling in the DS is mostly involved with reinforcement or reward learning and DRD2 is most commonly involved with aversion-formation (37). A positive correlation between *Drd1* expression and total ethanol consumption was observed in mock-colonized mice, but not in *C. albicans*-colonized mice suggesting dysregulation of the relationship of DRD1/reinforcement in these mice. Additionally, a negative correlation between *Drd2* expression and total ethanol consumption in *C. albicans*-colonized mice suggested that aversion to ethanol could develop. Therefore, there could be a lack of reinforcement learning, an active aversion, or a combination of both, that leads to lower ethanol preference in *C. albicans*-colonized mice. Results from the CTA suggest an aversion-dominated phenotype because colonized mice showed a more rapid acquisition of conditioned taste aversion. These findings support the conclusion that aversion to ethanol develops more readily in *C. albicans*-colonized mice.

When conducting other behavioral tests, we found that *C. albicans*-colonized mice injected with ethanol showed increased LORR onset, decreased locomotion in the open field test (OFT), and further impaired performance on the balance beam. The performance on the balance beam could be restored to mock-colonized performance by EP1/EP2 antagonists. These results suggest that PGE_2_ impact on EP1 and EP2 receptors could contribute to motor control following ethanol intoxication. The depressant and sedative effects of ethanol detected in the OFT and LORR tests could be driven by other systems impacted by *C. albicans* colonization such as the endocannabinoid system (ECS) (25). Both therapeutic manipulation and gene knockouts of the ECS modulated LORR and OFT phenotypes (54–56). Effects of *C. albicans* colonization on this pathway (25) could play a role in these phenotypes.

All of the experiments described in this communication were done with ethanol naïve mice. The effects of *C. albicans* colonization on mice chronically exposed to ethanol are currently unknown. There could be an altered effect of *C. albicans*-colonization on the host in the setting of AUD and/or chronic stress. Low DRD2 receptor abundance has been shown in individuals with AUD and predisposes mice to stress-induced increases in ethanol consumption (57,58). If DRD2 receptor abundance is reduced in humans with *C. albicans* colonization, as it is in colonized mice, these individuals may consume more ethanol under stress, possibly exacerbating ethanol consumption in AUD. However, our study focused only on initial alcohol consumption and these effects of *C. albicans* colonization in individuals with AUD and/or chronic stress are currently unstudied. This remains an area of important future research.

In summary, we have shown that GI colonization by *C. albicans* elevates serum PGE_2_ and these changes correlate with changes in *Ep* and *Drd* receptor expression in the dorsal striatum. *C. albicans* colonization decreased ethanol consumption and preference in C57BL/6 mice. These decreases in ethanol consumption correlated with changes in dopamine receptor expression and *C. albicans*-colonized mice showed altered behavioral effects of ethanol. In conclusion, we show for the first time, that changes in the fungal microbiome can impact host ethanol consumption and behavioral responses to ethanol.

## Materials and Methods

### Animals

Five-week-old female or male C57BL/6 mice (Jackson Laboratory) were acclimated to the facility and single-housing for 4-5 days with scruffing to acclimate to handling. Mice were orally inoculated with *C. albicans* at different times depending on the experiment. Mice were inoculated by pipetting 20uL containing 5x10^7^ cells of *C. albicans* strain CKY101(59) in phosphate buffered saline (PBS), or PBS only for mock-colonized mice, directly into the mouth of the mouse as previously described (25). Mock-colonized mice were given 20uL of PBS directly into their mouths. Mice were monitored throughout the experiment for colonization as previously described (25). No fungal colonies were observed in mock-colonized mice.

Mouse weight was measured throughout the experiment. Rarely, if a mouse lost 20% of its starting body weight, the mouse was euthanized. On the last day of each experiment, mice were anesthetized using the isoflurane drop method and euthanized via decapitation. Blood for serum and various organs were harvested for analysis. All mouse experiments were approved by the Tufts IACUC and were performed in accordance with IACUC guidelines for care and use of laboratory animals.

### Chemicals and Reagents

The EP1 antagonist SC-51089 (Cayman Chemical) and the EP2 antagonist TG6-10-1 (Cayman Chemical) were resuspended in 2:3:5 mixture of DMSO:sterile water:PEG400. Antagonists were resuspended and mixed at a concentration that would yield 2mg/kg of each antagonist in 50uL and 50 uL of the mixture was injected intraperitoneally.

Fifteen percent ethanol was prepared for consumption by diluting 95% ethanol (Fisher) with tap water. This solution was filter sterilized and added to sipper tubes for the two-bottle choice experiment.

16,16-dimethyl Prostaglandin E_2_ (Cayman Chemical) was dried under a gentle stream of nitrogen to evaporate the solvent. It was then resuspended in DMSO and injected into mice subcutaneously at a concentration of 25, 50, 100, or 300 ug/kg.

### Two-Bottle Choice Ethanol or Saccharin Consumption

During acclimation to the facility and single housing, mice also began acclimating to the sipper tubes used in the two-bottle choice experiment. Briefly, sterile acidified water was poured into 50 mL conical tubes (USA Scientific) and a sterile stopper (#6 Rubber, Ancare) with a sipper spout (Ancare) was placed into the 50 mL conical tube and wrapped with parafilm. Two of these sipper bottles with water were placed into standard cages for acclimation.

Following acclimation, mice were weighed and assigned to groups with equal weights. Mice were inoculated with *C. albicans* or mock-inoculated on day -1. On day 0, 15% (vol/vol) ethanol or 0.0125% (wt/vol) saccharin replaced one water bottle. Mouse weight and tube weight were then measured daily throughout the experiment. Sipper bottles were rotated daily to minimize side preference. After 2-4 days of ethanol consumption, mice were euthanized as previously described.

For ethanol preference, results show 26 mice per group from 3 replicate experimental trials. For saccharin preference, 14-15 mice per group from 2 experimental trials are shown.

### PGE-m ELISA

Trunk blood from mice was obtained after decapitation, collected in BD blood collection tubes (365967, Fisher). Tubes were then centrifuged in an Eppendorf table-top microcentrifuge at 10,000g for 10 minutes and serum was stored at -80°C. Serum was assayed with a Prostaglandin E Metabolite ELISA Kit (514531, Cayman) after removal of proteins by acetone precipitation. PGE-m ELISA was performed using the standard protocol, with derivatization, detailed in the product manual. Serum was diluted 15-25 times for the assay and OD was measured on a Biotek Epoch2 plate reader. A four parameter logistic curve was determined and raw PGE-m values were determined using the 4PL calculator by AAT Bioquest (60).

### Brain Dissection, RT-qPCR, and Primers

Mouse brains were dissected, flash frozen on aluminum foil chilled with dry ice and kept at -80°C. A brain matrix (ZIVIC Instruments, 5325) was allowed to chill on ice for 30 minutes. 2 mm sections of the brain from directly anterior of the cerebellum to the frontal pole were then cut using clean straight razor blades and the brain matrix. Sections were then observed to find landmarks indicating the location of the dorsal striatum as previously described (61). The dorsal striatum was dissected and RNA was extracted as previously described (25).

cDNA was then synthesized using random hexamer primers and RT-qPCR was performed as previously described (25). Primer sequences are listed in Supplemental Table S1.

### Ethanol-Induced Conditioned Taste Aversion

Following arrival, mice were acclimated to single-housing and sipper tubes by placing one sipper tube of water in their cages. Mice were weighed every other day to ensure they were consuming enough water to maintain their weight. The following protocol was adapted from a previous protocol that showed significant conditioned taste aversion in C57BL/6 mice (62). The first four days mice were acclimated to sipper tubes and single housing. Then, mice were subjected to 5 days of water restriction with only two hours of access to water for two hours daily (termed days 1-5 of the experiment). On day 5 of water restriction, mice were inoculated with *C. albicans* or PBS for mock-colonization and were re-inoculated every two days (days 5,7,9,11). On day 6, mice were given access to 1.2% saline for 1 hour in their home cages and injected with sterile saline, 2g/kg of ethanol, or 3g/kg of ethanol (20% vol/vol) immediately after the end of the 1-hour period. Five hours later, mice were given access to water for 30 minutes to rehydrate. This process was repeated for days 8 and 10; on day 12, mice were given access to 1.2% saline for 1 hour in their home cages and then euthanized. Fecal pellets were collected for CFU determination on days 6, 8, 10, and cecum contents were collected on day 12. Data were analyzed as a percentage of saline consumption compared to the initial amount consumed on day 6.

### Open Field Activity Test

On the open field test (OFT) habituation day, mice were injected an average of 15 minutes before their respective trials with 100uL of sterile saline intraperitoneally with an insulin syringe. Following 15 minutes, mice were gently placed in the center of a 37cm X 37cm plastic open field box. Mice were allowed to roam freely for 10 minutes and removed from the apparatus. The apparatus was then sterilized using 70% ethanol. This protocol was repeated for every mouse.

On OFT test day, mice were injected with 1.5g/kg of ethanol or sterile saline intraperitoneally an average of 15 minutes before their trials. The volume of injection varied from approximately 90-110uL depending on the weight of the mouse. The ethanol concentration was adjusted so that a mouse of average weight for the group would be injected with 100uL ethanol solution. The 10-minute test run and cleaning of the apparatus were repeated as above for every mouse. Mouse behavior was recorded using a web camera. Scoring videos was performed using EthovisionXT software to automatically determine distance traveled. Total distance traveled was plotted relative to the mean of the sterile saline group distance traveled.

### Balance Beam

A balance beam apparatus was constructed using a square 0.25-inch diameter, 3-foot-long metal rod held approximately 1.5ft above the surface. A plastic box to hold bedding was affixed to one end to provide incentive for the mouse to cross the entirety of the balance beam. A reading lamp shining on the start zone of the beam was used as an aversive stimulus. Start and end points for the balance beam trials spanning 65 cm of the beam were marked.

On habituation day, mice were injected with 100uL of sterile saline intraperitoneally approximately 15 minutes before the start of their first of 3 trials. Mice were then placed in the start zone to begin trial 1. After completing the trial, mice were given 30 seconds of rest in their home cage before the start of trial 2. Trials 2 and 3 were performed by the same procedure. After a mouse completed 3 trials, the balance beam was cleaned using 70% ethanol and the next mouse was tested. In the event that a mouse would freeze, or very rarely fall, they would be gently prodded or placed back on the bar until they completed the trial. All mice within a cohort underwent this procedure.

On test day, mice were injected with 100uL of sterile saline or injected with 1.5g/kg of ethanol as described in the **open field activity test** section. These injections were performed approximately 15 minutes before the start of the first trial. A web camera was used to record mouse behavior during these trials. The 3 trials were repeated as described above. All mice completed the entirety of the 3 trials.

Videos were blinded for scoring. Start and stop times were determined when the mouse’s nose crossed designated marks on the bar. A slip was determined when a mouse’s foot slipped below the bar or a significant slip from a standing position on the bar to the pelvis touching the bar occurred. A fall was scored when the mouse slipped to the underside of the bar. Very rarely a mouse would fall from the bar into safety padding below the bar. Both of these events were scored equally as falls. Time or slips and falls were summed for all 3 trials for each mouse.

### Loss of Righting Reflex

Mice were co-housed with another mouse within the same group for the LORR. Their nestlet was removed for the test. Mice were injected with 3.5g/kg of ethanol intraperitoneally and the time of injection was recorded. Within roughly 1-2 minutes most mice lost righting reflex which was defined as an inability to right themselves when lying on their back in a V-shaped plastic trough. The time when a mouse lost righting reflex was designated the LORR onset. Six minutes after injection, mice were given one minute of exposure to isoflurane via the drop method and euthanized as above. Occasionally, a mouse would have to be removed from the data due to shearing of an artery during injection—significantly altering LORR time.

### Blood Ethanol Concentration Assay

Following LORR, trunk blood was collected and serum was extracted as previously described in the **PGE-m ELISA** section. For the blood ethanol concentration determination, an EnzyChrom Ethanol Assay Kit (ECET-100, BioAssay Systems) was used. Standard protocols were followed for the determination using serum that was diluted 15X. OD was measured on a Biotek Epoch2 plate reader.

## Acknowledgements

We thank Dr. Jesus Romo, Dr. Paola Zucchi, Dr. Ashlee Junier, Andressa Barossi Pesarini, Dr. Laura Markey, and Adrianne Gladden-Young for helpful discussion. We also thank Nick D’Arrigo and Emma Hayes for technical assistance.

## Supplemental Figures

**Figure S1:**
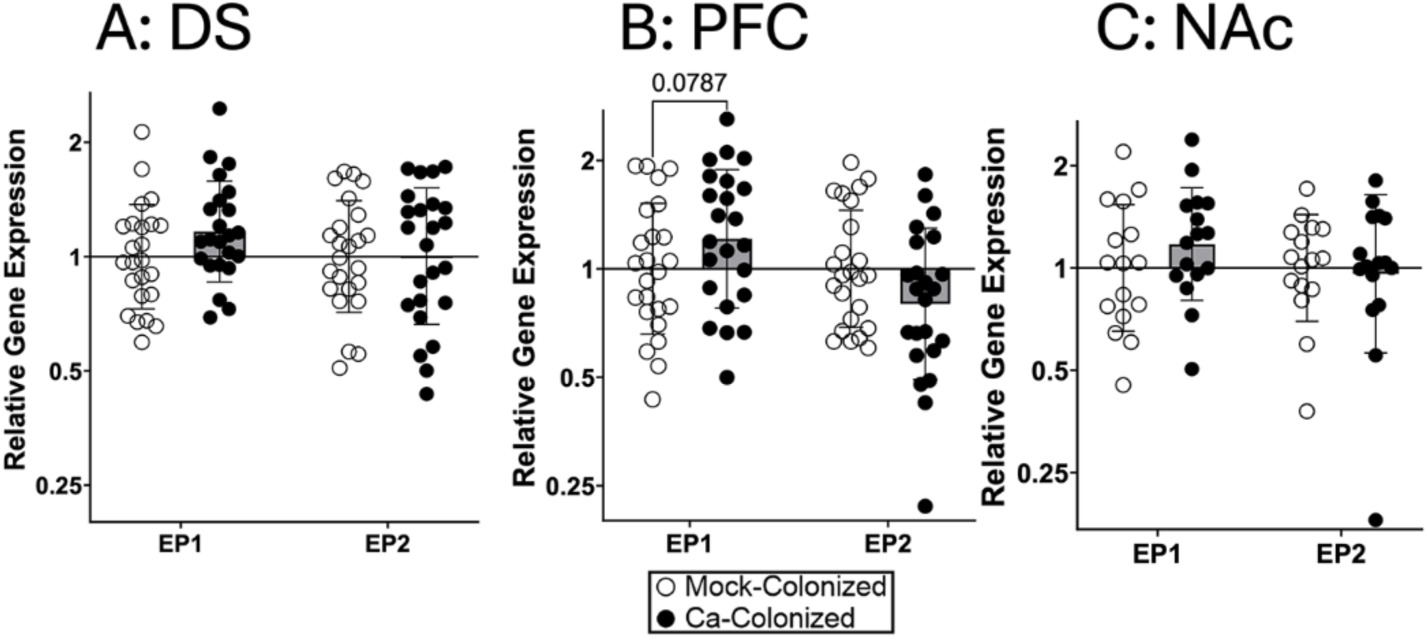
Ep receptor expression were not different in C. albicans-colonized mice vs mock-colonized mice in various brain regions. Single housed female C57BL/6 mice were orally inoculated with *C. albicans* strain CKY101 or PBS and subjected to the 2-bottle choice experiment as in Figure 1. On day 2, mice were euthanized, and brains were collected. *Ep* receptor expression was measured in various brain regions of mock-colonized or *C. albicans*-colonized mice by RT-qPCR using the ddCT method. Geometric mean and geometric standard deviation are shown. A Two-way ANOVA was performed for statistics and p-values are shown. **(A)** Shows expression in dorsal striatum (DS) **(B)** Shows expression in the prefrontal cortex (PFC) **(C)** Shows expression in the nucleus accumbens (NAc)

**Figure S2:**
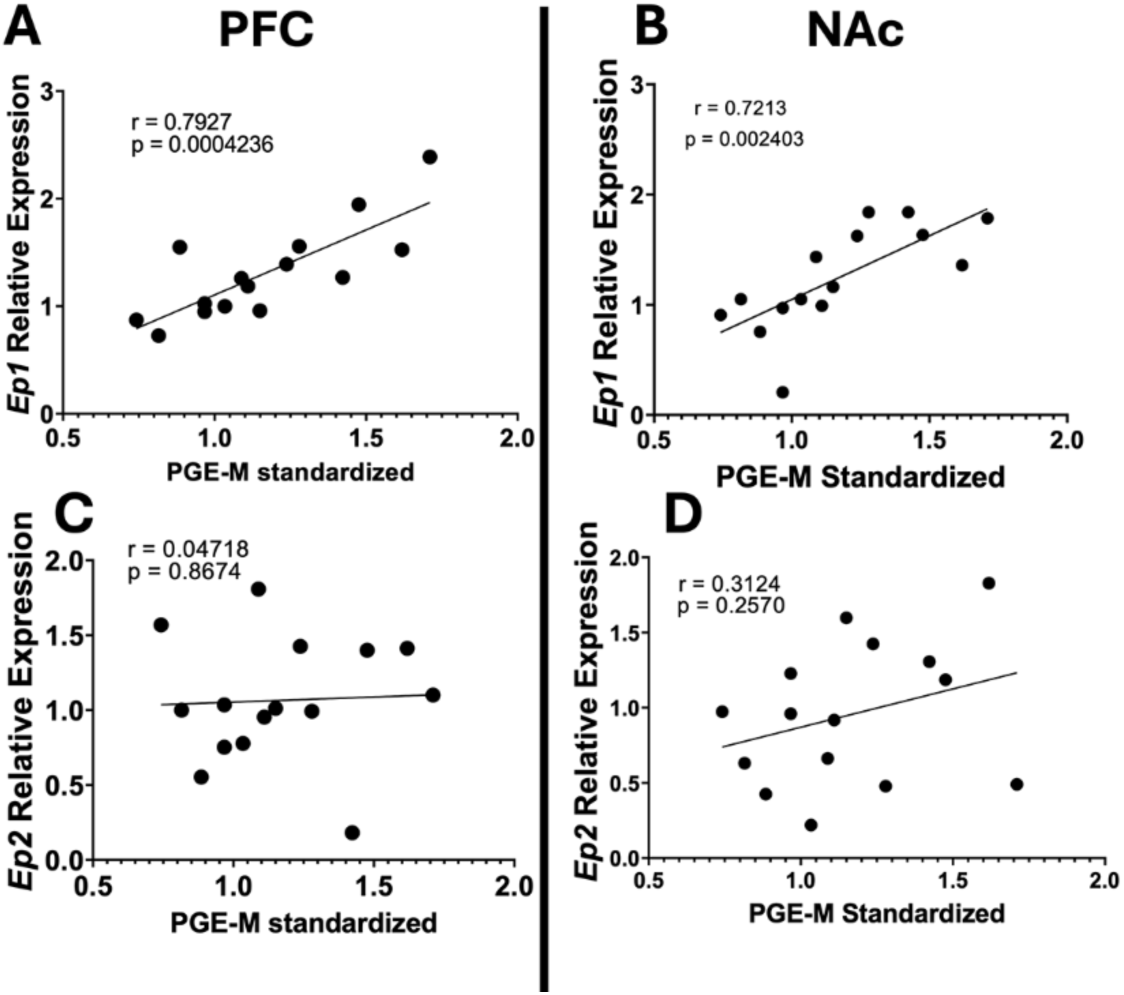
Serum PGE-metabolite correlates with EP1 Receptor expression in the PFC and NAc of C. albicans-colonized mice. Single housed female C57BL/6 mice were orally inoculated with *C. albicans* strain CKY101 or PBS and subjected to the 2-bottle choice experiment as in Figure 1. On day 2, mice were euthanized, and brain and serum were collected. Expression of *Ep* genes relative to average expression in mock-colonized mice from the same experimental trial, plotted as a function of concentration of PGE-metabolite standardized to the average in mock-colonized mice in the same experimental trial. Each symbol represents an individual mouse. **(A,C)** show gene expression of *Ep1* and *Ep2*, respectively, in the prefrontal cortex (PFC). **(B,D)** show gene expression of *Ep1* and *Ep2*, respectively, in the nucleus accumbens (NAc). Pearson correlations were used to test for significant correlations, r=correlation strength and p=statistical significance. Significant correlations were observed between *Ep1* expression and concentration of PGE-M in the PFC and NAc of *C. albicans-*colonized mice.

**Figure S3:**
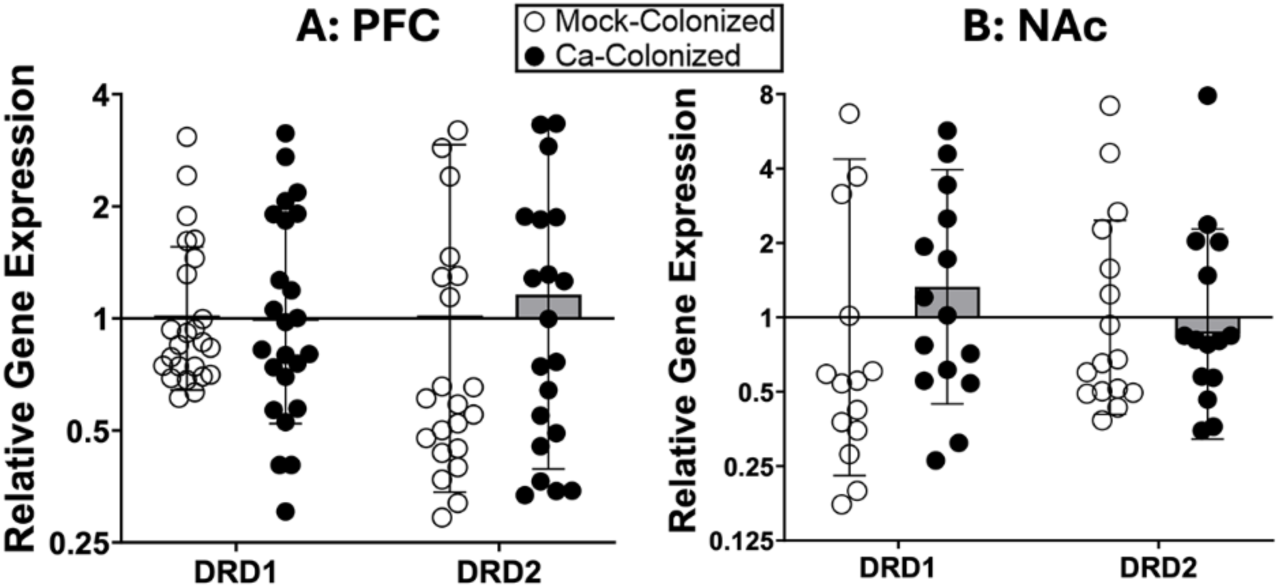
Drd gene expression was not different between C. albicans-colonized and mock-colonized mice in the PFC and NAc. Single housed female C57BL/6 mice were orally inoculated with *C. albicans* strain CKY101 or PBS and subjected to the 2-bottle choice experiment as in Figure 1. On day 2, mice were euthanized, and brains were collected. *Drd* receptor expression was measured in the prefrontal cortex (PFC) **(A)** or nucleus accumbens (NAc) **(B)** of mock-colonized or *C. albicans*-colonized mice by RT-qPCR using the ddCT method. Geometric mean and geometric standard deviation are shown. Multiple t-tests performed for statistics—no significant differences were observed.

**Figure S4:**
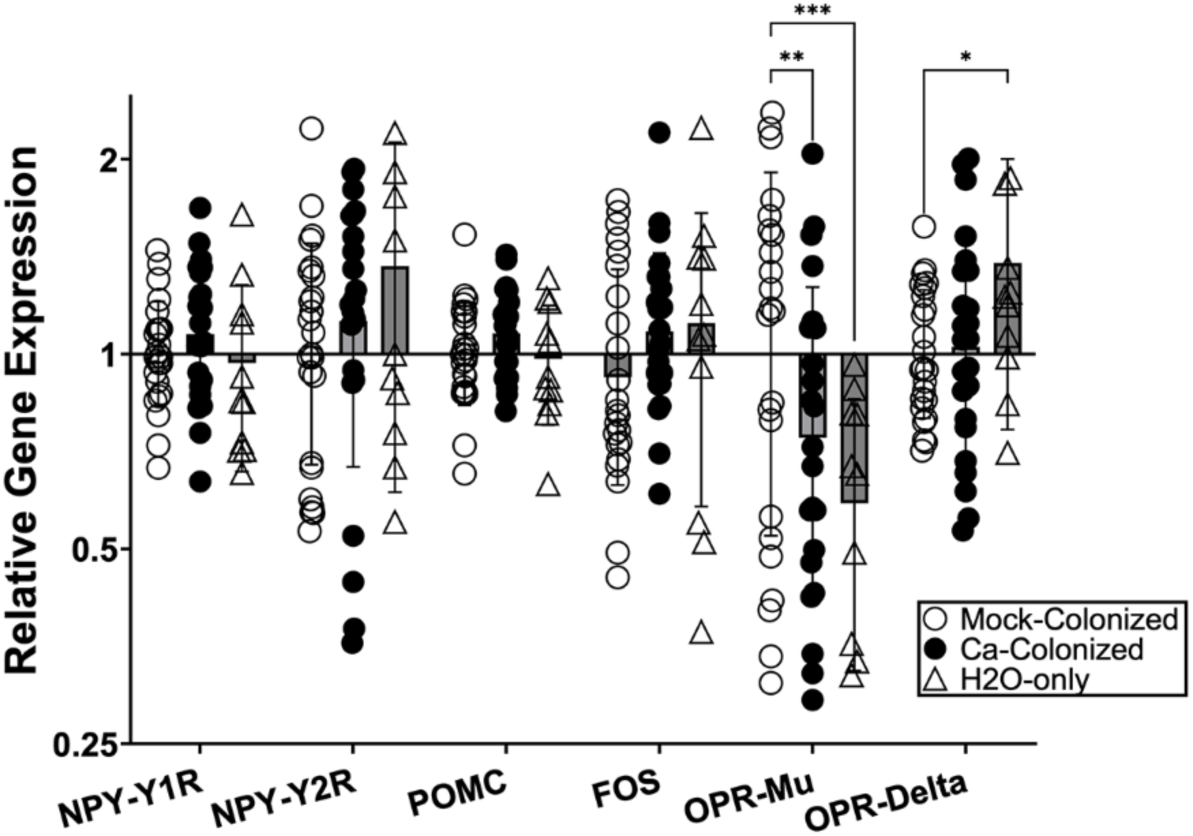
Expression of other addiction-implicated genes in the Dorsal Striatum of C. albicans-colonized mice. Single housed female C57BL/6 mice were orally inoculated with *C. albicans* strain CKY101 or PBS and subjected to the 2-bottle choice experiment as in Figure 1. On day 2, mice were euthanized, and brains were collected. Groups of mice that consumed ethanol (Ca-colonized or Mock-colonized) or mice that were mock-colonized and only given access to water (H2O-only) are shown. The expression of genes implicated in addiction was measured in the dorsal striatum of the different groups by RT-qPCR using the ddCT method. Geometric mean and geometric standard deviation are shown. A two-way ANOVA with Dunnett’s correction was performed for statistics. * p=0.0315; ** p=0.0065; *** p=0.0001.

**Figure S5:**
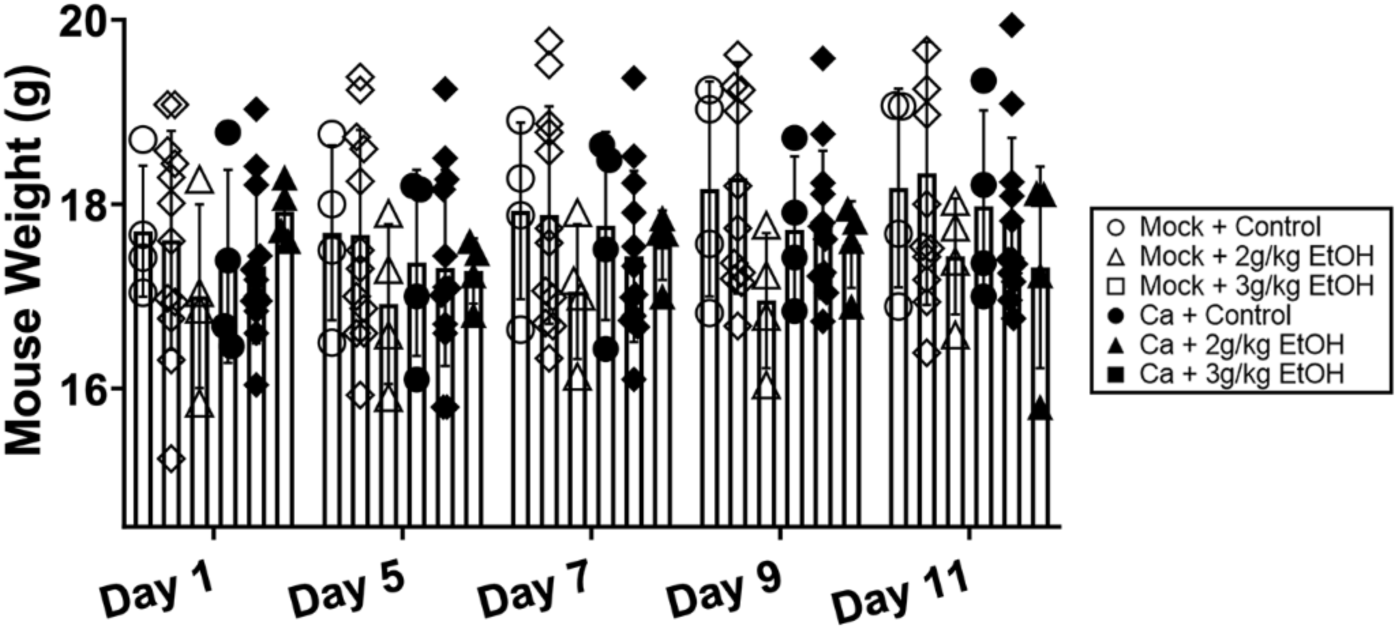
No differences in mouse weight were observed in the ethanol-induced conditioned taste aversion experiment. Single housed female C57BL/6 mice were trained to drink their daily liquid in the late morning. Mice orally inoculated with *C. albicans* strain CKY101 or PBS were given one hour of access to a novel tastant, 1.2% saline solution, and then injected intraperitoneally with ethanol (2 g/kg or 3 g/kg) or control (sterile saline). Mouse weight is shown in grams. Mock-colonized, control (open circles); mock-colonized + 2g/kg EtOH (open triangles); mock-colonized + 3 g/kg EtOH (open squares); *C. albicans*-colonized, control (closed circles); *C. albicans-*colonized +2 g/kg EtOH (black triangles); *C. albicans-*colonized + 3 g/kg EtOH (closed squares). Bars show means, error bars show standard deviation, and each symbol represents one mouse. A two-way ANOVA corrected for repeated measures was performed for statistical significance—there were no significant comparisons.

**Figure S6:**
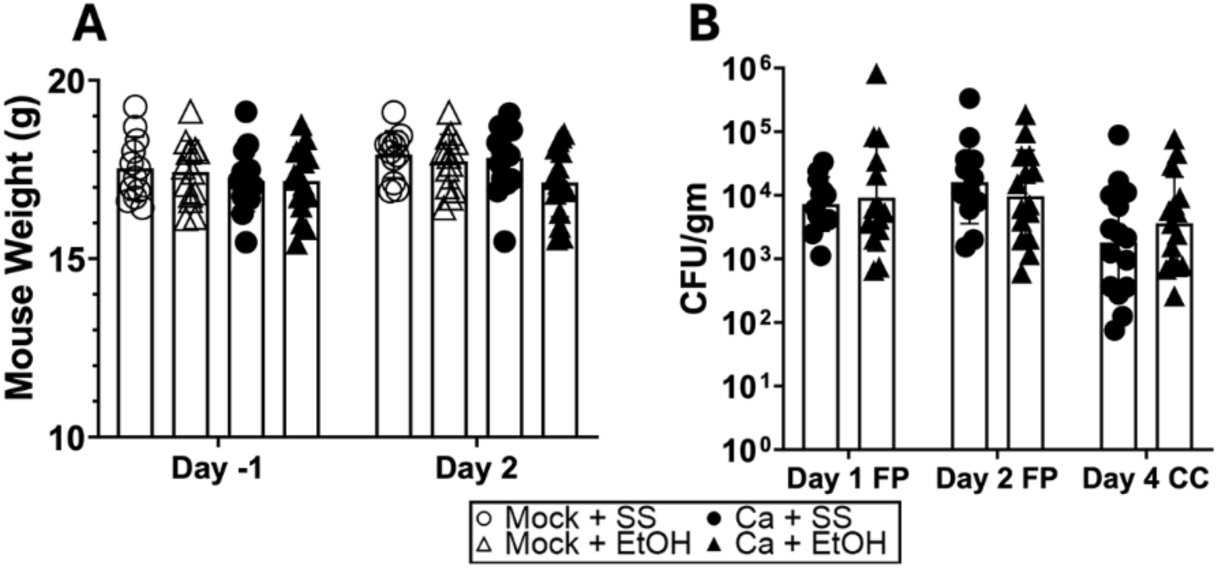
No differences in mouse weight or colonization were observed in the LORR, OFT, and balance beam experiments. Single housed female C57BL/6 mice were orally inoculated with *C. albicans* strain CKY101 or PBS on days 0 and 2, and subjected to the behavioral tests illustrated in Fig. 8A**. (A)** Mouse weights on days -1 and 2 are shown. **(B)** CFU/gm fecal pellets (FP) or cecum contents (CC) collected on days indicated. Bars show means with the standard deviation (A) or geometric mean with geometric standard deviation (B). Symbols represent individual mice. Two-way ANOVAs corrected for multiple repeated measures were performed for statistics and there were no significant differences.

**Figure S7:**
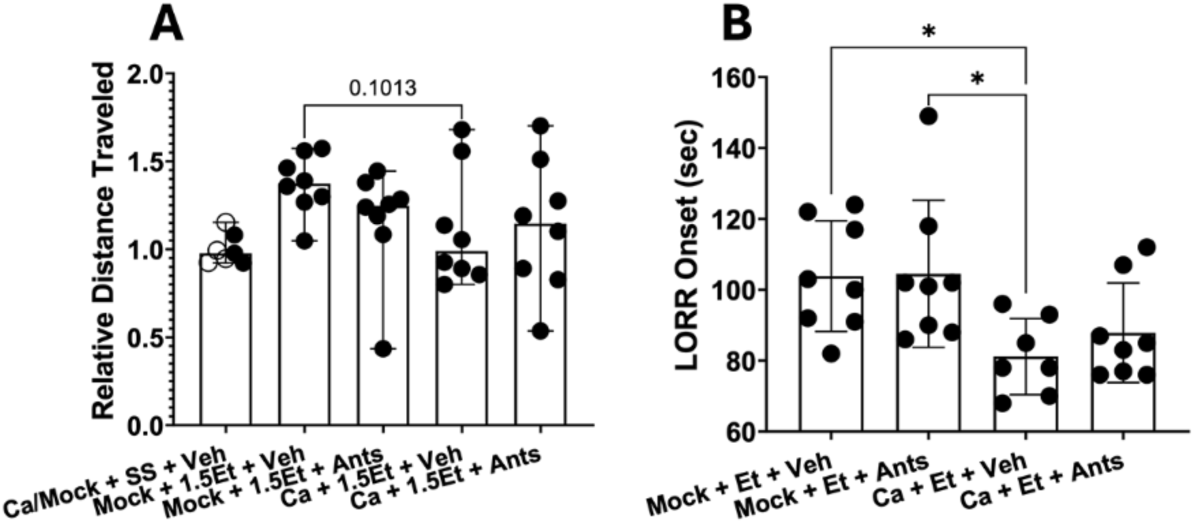
Antagonism of Ep receptors did not change the behavior of mice in the OFT or LORR tests. Single housed female C57BL/6 mice were orally inoculated with *C. albicans* strain CKY101 or PBS on days 0 and 2, and subjected to the behavioral tests illustrated in Figure 8A, except that mice were injected intraperitoneally with EP1 and EP2 antagonists or vehicle daily on days 0-4 approximately one hour before each behavioral test. **(A)** Groups are defined as: Ca/Mock + SS + Veh: mock-colonized or *C. albicans-*colonized, received sterile saline ip injections on days 1 and 3 and vehicle ip injections daily; Mock + EtOH + Veh: mock-colonized, received 1.5g/kg ethanol ip injected on days 1 and 3 and vehicle ip injections daily; Mock + EtOH + Ants: mock-colonized, received 1.5g/kg ethanol ip injected on days 1 and 3 and antagonist ip injections daily; Ca + EtOH + Veh: *C. albicans*-colonized, received 1.5g/kg ethanol ip injected on days 1 and 3 and vehicle ip injections daily; Ca + EtOH + Ants: *C. albicans*-colonized, received 1.5g/kg ethanol ip injected on days 1 and 3 and antagonist ip injections daily **(B)** Groups are defined as: Mock + Et + Veh: mock-colonized, received 1.5g/kg ethanol ip injected on days 1 and 3 and 3.5g/kg EtOH on day 4 and vehicle daily; Mock + Et + Ants: mock-colonized, received 1.5g/kg ethanol ip injected on days 1 and 3 and 3.5g/kg EtOH on day 4 and antagonists daily; Ca + Et + Veh: *C. albicans*-colonized, received 1.5g/kg ethanol ip injected on days 1 and 3 and 3.5g/kg EtOH on day 4 and vehicle daily; Ca + Et + Ants: *C. albicans*-colonized, received 1.5g/kg ethanol ip injected on days 1 and 3 and 3.5g/kg EtOH on day 4 and antagonists daily. **(A)** distance traveled in the open field test in a ten-minute trial relative to the average for sterile saline-injected (SS) mice within the same group (mock-colonized or *C. albicans*-colonized). A Brown-Forsythe and Welch ANOVA test was run for statistical significance. **(B)** time after injection (seconds) for the mouse to lose its righting reflex (LORR onset). An ordinary One-way ANOVA was completed for statistical significance. **\*** p<0.0465.

**Table S1:**
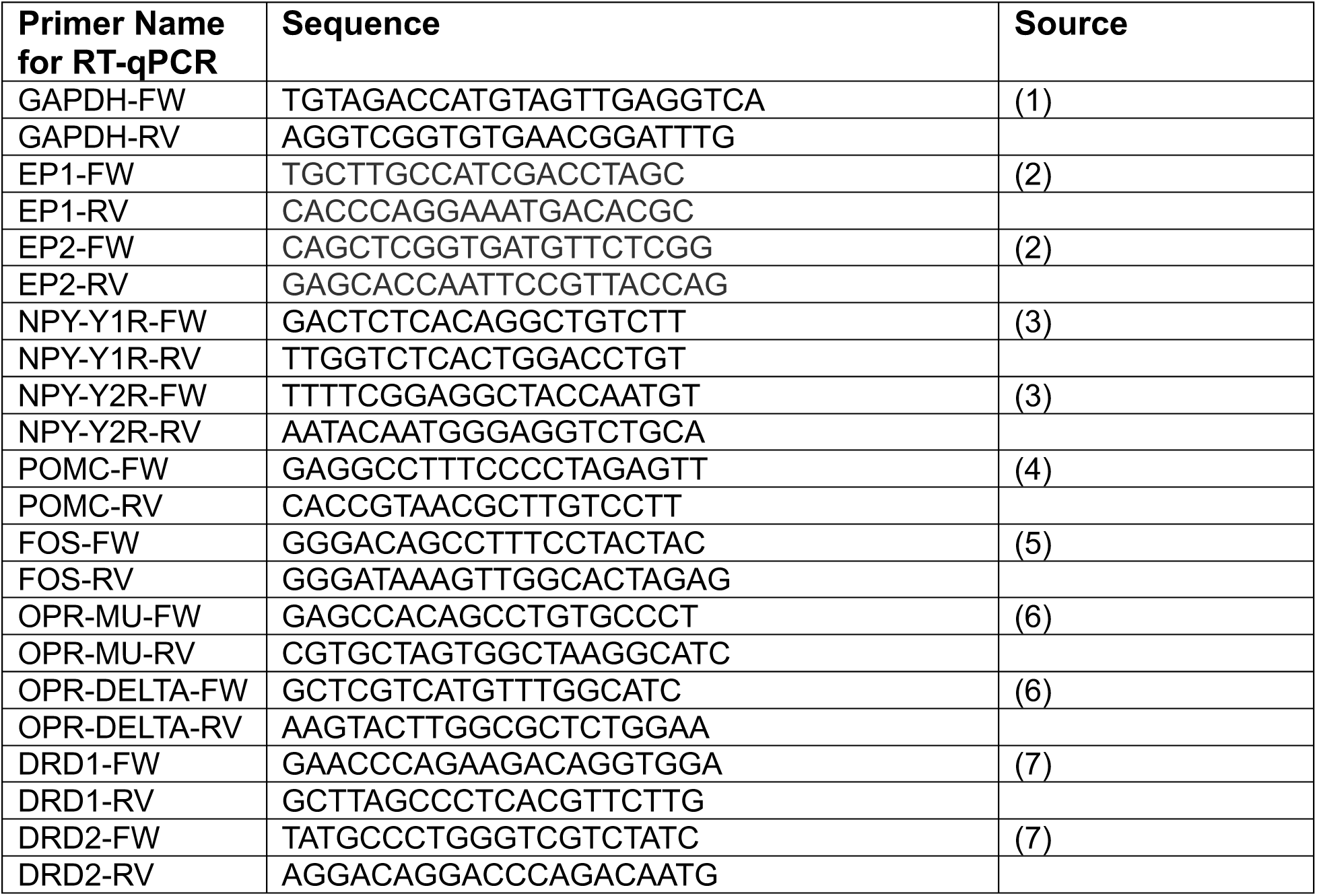
Primers used In this study.

